# Prediction and Characterization of RXLR Effectors in *Pythium* Species

**DOI:** 10.1101/2019.12.19.882209

**Authors:** Gan Ai, Kun Yang, Yuee Tian, Wenwu Ye, Hai Zhu, Tianli Li, Yaxin Du, Qingyue Xia, Danyu Shen, Maofeng Jing, Ai Xia, Daolong Dou

## Abstract

Being widely existed in oomycetes, the RXLR effector features conserved RXLR-dEER motifs in its N terminal. Every known *Phytophthora* or *Hyaloperonospora* pathogen harbors hundreds of RXLRs. In *Pythium* species, however, none of the RXLR effectors has been characterized yet. Here, we developed a stringent method for *de novo* identification of RXLRs and characterized 359 putative RXLR effectors from nine tested *Pythium* species. Phylogenetic analysis revealed a single superfamily formed by all oomycetous RXLRs, suggesting they descent from a common ancestor. RXLR effectors from *Pythium* and *Phytophthora* species exhibited similar sequence features, protein structures and genome locations. In particular, the mosquito biological agent *P. guiyangense* contains a significantly larger RXLR repertoire than the other eight *Pythium* species examined, which may result from gene duplication and genome rearrangement events as indicated by synteny analysis. Expression pattern analysis of RXLR-encoding genes in the plant pathogen *P. ultimum* detected transcripts from the vast majority of predicted *RXLRs* with some of them being induced at infection stages. One such RXLRs showed necrosis-inducing activity. Furthermore, all predicted *RXLRs* were cloned from two biocontrol agents *P. oligandrum* and *P. periplocum*. Three of them were found to encode effectors inducing defense response in *Nicotiana benthamiana*. Taken together, our findings represent the first complete synopsis of *Pythium* RXLR effectors, which provides critical clues on their evolutionary patterns as well as the mechanisms of their interactions with diverse hosts.

**Author summary:** Pathogens from the *Pythium* genus are widespread across multiple ecological niches. Most of them are soilborne plant pathogens whereas others cause infectious diseases in mammals. Some *Pythium* species can be used as biocontrol agents for plant diseases or mosquito management. Despite that phylogenetically close oomycete pathogens secrete RXLR effectors to enable infection, no RXLR protein was previously characterized in any *Pythium* species. Here we developed a stringent method to predict *Pythium* RXLR effectors and compared them with known RXLRs from other species. All oomycetous RXLRs form a huge superfamily, which indicates they may share a common ancestor. Our sequence analysis results suggest that the expansion of RXLR repertoire results from gene duplication and genome recombination events. We further demonstrated that most predicted *Pythium RXLRs* can be transcribed and some of them encode effectors exhibiting pathogenic or defense-inducing activities. This work expands our understanding of RXLR evolution in oomycetes in general, and provides novel insights into the molecular interactions between *Pythium* pathogens and their diverse hosts.

## Introduction

Coevolution of microbial pathogens and host plants is driven by their endless arms race [1]. Plants respond to the conserved pathogen-associated molecular patterns (PAMPs) of pathogens via a diverse group of cell surface receptors, and thereby induce PAMP-triggered immunity (PTI) [2]. Numerous plant pathogens, including bacteria, nematodes, fungi and oomycetes, can counteract PTI by delivering effector proteins into host cells to suppress plant defense and facilitate infection [1,3].

RXLR proteins are a group of effectors initially identified from *Phytophthora* species. Their N terminals feature conserved RXLR-dEER motifs, which is assumed to function in the host cell translocation process. In contrast, the C-terminals of RXLR proteins are relatively divergent due to their distinct effector activities in modulating host immunity [4,5,6]. The interacting targets of some *Phytophthora* and *Hyaloperonospora* RXLRs have been well-studied. For example, the RXLR effector PexRD2 from *Phytophthora infestans* can perturb host resistance response by interacting with a positive plant immunity regulator MAPKKK [7]. PsAvh262, which is essential for the pathogenicity of *Phytophthora sojae*, suppresses ER stress-dependent immunity via stabilizing ER-luminal binding immunoglobulin proteins (BiPs) [8]. A *Phytophthora capsici* RXLR effector, PcRXLR207, triggers the degradation of BPA1 (Binding partner of ACD11) family proteins to facilitate the biotroph to necrotroph transition [9]. A conserved RXLR effector HaRxL23 from *Hyaloperonospora arabidopsidis* can suppress PTI in tobacco as well as effector-triggered immunity (ETI) in soybean [10]. Furthermore, some RXLRs are avirulence proteins exhibiting gene-for-gene interactions with specific host resistance proteins [11].

Quantities of RXLR candidates have been identified from *Hyaloperonospora* and *Phytophthora* species via bioinformatic searching approaches. A pilot method was designed to search for protein with a signal peptide (SP) in its first 30 residues and an RXLR motif within the following 30 residues. This method was used for RXLR identification in *Hyaloperonospora parasitica*, *Phytophthora ramorum* and *P. sojae* [12]. Later on, RXLRs were identified in three *Phytophthora* species using the hidden Markov model (HMM) together with regular expression (regex) model [5]. A homology searching method was also developed and used together with HMM to predict 370 and 392 RXLRs in *P. ramorum* and *P. sojae*, respectively [13]. Moreover, the three approaches above were combined to characterize RXLRs in *Plasmopara viticola,* an oomycete pathogen infecting grapevine [14]. RXLR genes and proteins predicted from different species show similar features of diverse variation, disorder region enrichment and predominant localization in gene-sparse regions [13,15,16]. Interestingly, there are also putative RXLRs found in *Albugo* and *Saprolegnia* species. For example, 26 Ac-RxL effectors were identified in *Albugo candida* by searching for the RXL string within proteins with SP. These Ac-RxLs exhibit effector activity despite that they only contain a degenerate RXLR motif [17]. In *Albugo laibachii,* 25 RXLR and 24 RXLQ effector candidates were found using similar methods with two of them being functionally validated [18]. The fish-infecting oomycete *Saprolegnia parasitica* harbors an RXLR effector that enters the host cells to promote virulence [19]. *Pythium* belongs to *Pythiales*, a sister order of *Peronosporales, Albuginales* and *Saprolegniales* [20]. All these RXL-containing oomycetes are phylogenetically close to *Pythium* species. Nevertheless, previously reported *Pythium* genome analyses did not reveal any RXLR-encoding genes [21,22,23,24].

Unlike *Phytophthora* species which are mostly plant-infecting pathogens, *Pythium* species occupy multiple ecological niches. Many *Pythium* species, including *P. ultmum, P. iwayamai*, and *P. aphanidermatum*, cause a wide variety of diseases in plants [21,22], whereas *P. insidiosum* is a notorious pathogen infecting human and animals [23]. Interestingly, Some *Pythium* species, such as *P. oligandrum* and *P. periplocum*, are used as mycoparasitic biocontrol agents since they can infect fungal hosts and induce plant defense response as well [25,26,27]. Likewise, *P. guiyangense* is a parasite used for mosquito control [24].

It is still unknown whether *Pythium* genomes encode RXLR effectors, or which kinds of effectors participate in the interactions between *Pythium* species and their broad hosts. To answer the questions above, we developed a stringent method for *de novo* identification of RXLRs in the draft genome sequences of 9 *Pythium* species and performed comprehensive genomic analysis on the RXLRs predicted. *Pythium* RXLRs form a single superfamily with all other known oomycetous RXLRs. All RXLR genes and proteins exhibit similar features in sequence divergence, disorder content and genome location. Four novel RXLRs from *P. ultimum, P. oligandrum* and *P. periplocum* were functionally verified as effectors. Our findings reveal the wide occurrence of RXLRs in *Pythium* species at the whole-genome level, demonstrate the effector activities of selected *Pythium* RXLRs, and provide a solid platform for investigating the diversified roles of *Pythium* RXLRs in pathogenicity.

## Results

### *De novo* identification of RXLR effectors

No *Pythium* RXLR effector has been previously identified despite the numerous reports of RXLRs in their phylogenetically close species [21,22,23,24]. Here we developed a *de novo* identification method with stringent threshold to screen RXLR candidates in 9 *Pythium* species (Fig 1A). Meanwhile, well-studied *P. sojae, P. ramorum* and *H. arabidopsidis* genomes were used as positive controls. Some other species including two diatoms were also parallelly examined (S1 Table, Fig 1B). For every gene, all 6 possible open reading frames (ORFs) were retrieved from each genome because almost all *RXLRs* lack intron [13]. Sequences were considered as sORFs (open reading frames encoding putative secretory protein) if their encoding proteins contain an SP but lack transmembrane region (TM).

**Fig 1.**
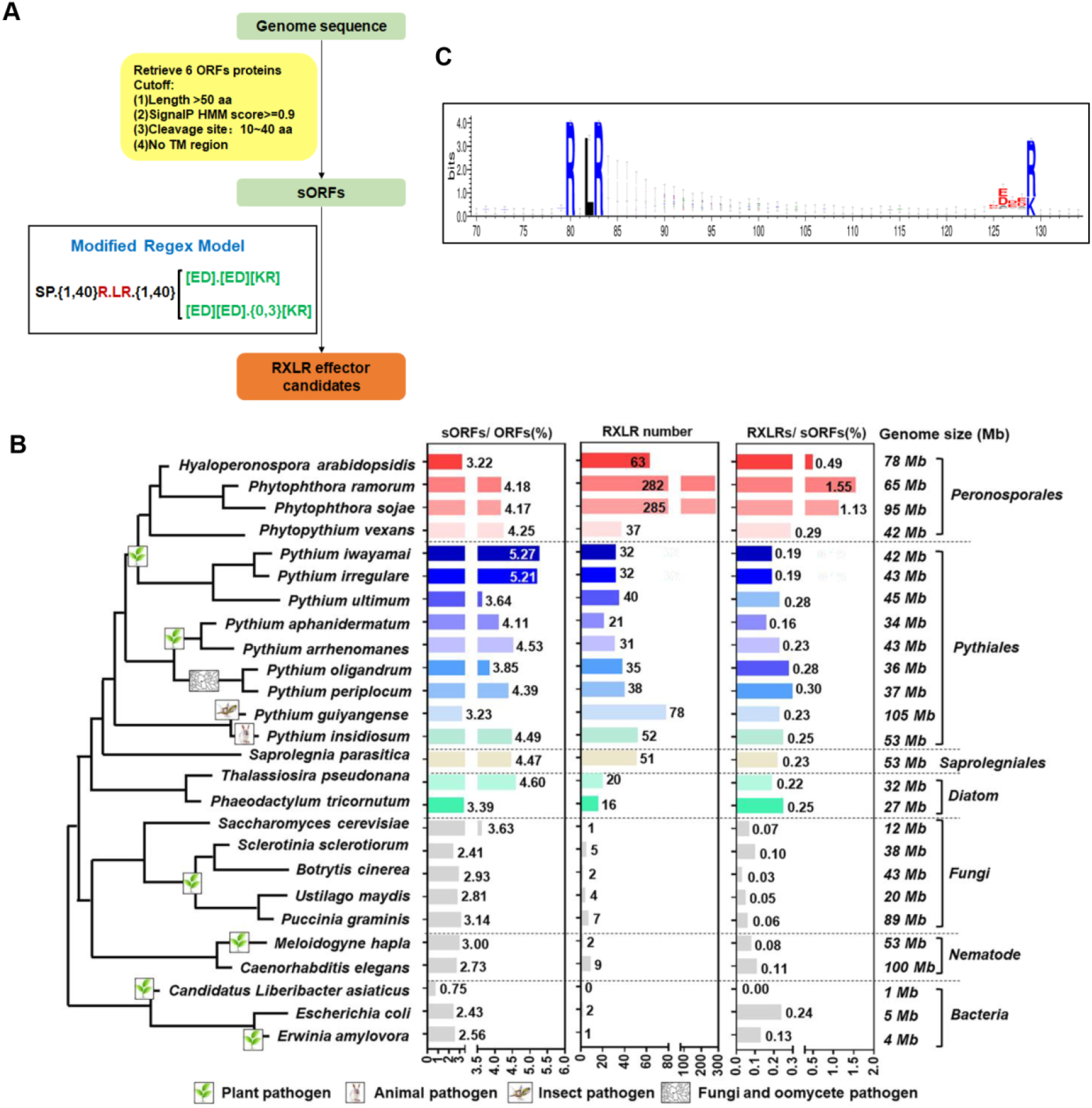
*De novo* identification of RXLR effectors in *Pythium* species. (A) The RXLR effector identification pipeline. sORFs indicate open reading frames encoding secretory proteins with a signal peptide (SP) but without transmembrane region (TM). (B) Summary of predicted RXLRs. Phylogeny of the 26 species is based on data from the Taxonomy Database and previous studies. The ratios of sORFs to whole ORFs, the counts of RXLR candidates and the ratios of RXLRs to sORFs in each species are showed in boxplots following the species names. The genome size of each species is showed after the corresponding boxplots. (C) Weblogo of the RXLR-dEER motifs of *Pythium* RXLR effectors.

In total, 311,164 sequences were achieved with 12,490 to 33,431 sORFs being found in each *Pythium* species, which is close to that of *Hyaloperonospora* and *Phytophthora* (12,830 to 25,178 sORFs). Interestingly, non-oomycetes only had 47 to 12,213 sORFs identified (S1 Table). Significantly more sORFs can be found in oomycetes when compared to other species (*P*<2.2E–16, Fisher’s Exact Test), indicating the occurrence of more expansion events in secretory protein-encoding genes in oomycetes.

Since some functionally verified RXLR effectors contain degenerate dEER motifs (S2 Table), the original regex model (SP.{1,96}R.LR.{1,40}[ED][ED][KR]) [5] was modified (SP.{1,40}R.LR.{1,40}([ED].[ED][KR]|[ED][ED].{0,3}[KR]) (Fig 1A) to allow the match of these atypical RXLRs. In total, 1,146 RXLR candidates were identified using this modified model (Fig 1B).

We found 63, 282 and 285 RXLRs in *H. arabidopsidis*, *P. ramorum* and *P. sojae*, respectively. They are all subsets of previously identified RXLRs and account for 42%, 53% and 42% of the total sets, respectively [5,12,13,28]. In contrast, only 0 to 9 RXLRs were predicted in fungi, nematodes or bacteria. These control results demonstrate the high reliability as well as the low false-positive rate of our method. 21 to 78 putative RXLRs were identified in each *Pythium* species. All predicted *Pythium* RXLR proteins contain a conserved SP, an RXLR motif and a dEER motif, in which two acidic amino acids (D or E) are enriched and followed by a conserved basic amino acid (R or K) (Fig 1C, S1 Fig).

To test the reliability of RXLR prediction in *Pythium*, an enrichment assay was conducted by comparing the number of RXLRs in each oomycete species with that of the fungal pathogen *Botrytis cinerea* (randomly selected as a control). All oomycetes including the *Pythium* species showed significant enrichment of RXLR sequences (*P*<0.01, Fisher’s Exact Test) (S3 Table). The enrichment was not observed in any of the non-*Stramenopiles* species (*P*>0.05, Fisher’s Exact Test). Next, a permutation test was performed to evaluate whether the predicted RXLR effectors are randomly occurred. We permuted the 90 after-SP residues of all sORFs and re-performed RXLR searching with the modified model for 100 times to estimate the false-positive rates. In general, very few RXLR predictions were generated from the permuted sequences, indicating that the vast majority of oomycete RXLRs were not predicted by chance (S4 Table). For example, 35 RXLRs were predicted in *P. oligandrum* whereas only 1 RXLR on average detected in its permuted sequences.

Besides their presence in all examined oomycetes including *Pythium* species, RXLR sequences can also be identified in diatoms (20 in *Thalassiosira pseudonana* and 16 in *Phaeodactylum tricornutum*) with significant enrichment (*P*<0.01, Fisher’s Exact Test). They may be the precursors of RXLR effectors that help diatoms compete with other organisms in the complicated environment.

### The evolutionary patterns of oomycete RXLRs

Despite the highly divergence of RXLR proteins resulted from host-pathogen arms race, RXLR effectors from *P. sojae* and *P. ramorum* exhibit significant relatedness, suggesting that the hundreds of fast-evolving RXLRs in these two species may derive from a single common ancestor [13]. Therefore, we used the BLASTP software to analyze whether the predicted *Pythium* RXLRs belong to the same evolutionary theme. The highly similar SP regions were removed to avoid their interference. Using BLASTP hit *E*-value<1 as the cutoff, 840 RXLR proteins had significant hits to one or more other RXLRs. In total, 15,071 RXLR pairs were identified, which form a huge relatedness network (Fig 2A). *Phytophthora* RXLRs have a higher within-genus relatedness level when compared with those from *Pythium* or *Hyaloperonospora*, suggesting a larger contribution of recent gene expansion events in *Phytophthora*. We found three clusters sharing little relatedness with *Phytophthora* RXLRs, which indicates their relative independence in evolution. Almost all RXLRs in these clusters are from *Pythium*. The three clusters are named as Cluster 1 (C1), Cluster 2 (C2) and Cluster 3 (C3), respectively (Fig 2A). For example, the average sequence identity of the 22 RXLRs in C1 is 54%. A Peptidase_S8 domain (PF00082.22), which is absent in *Phytophthora*, can be found in the C-terminal of 21 C1 RXLRs.

**Fig 2.**
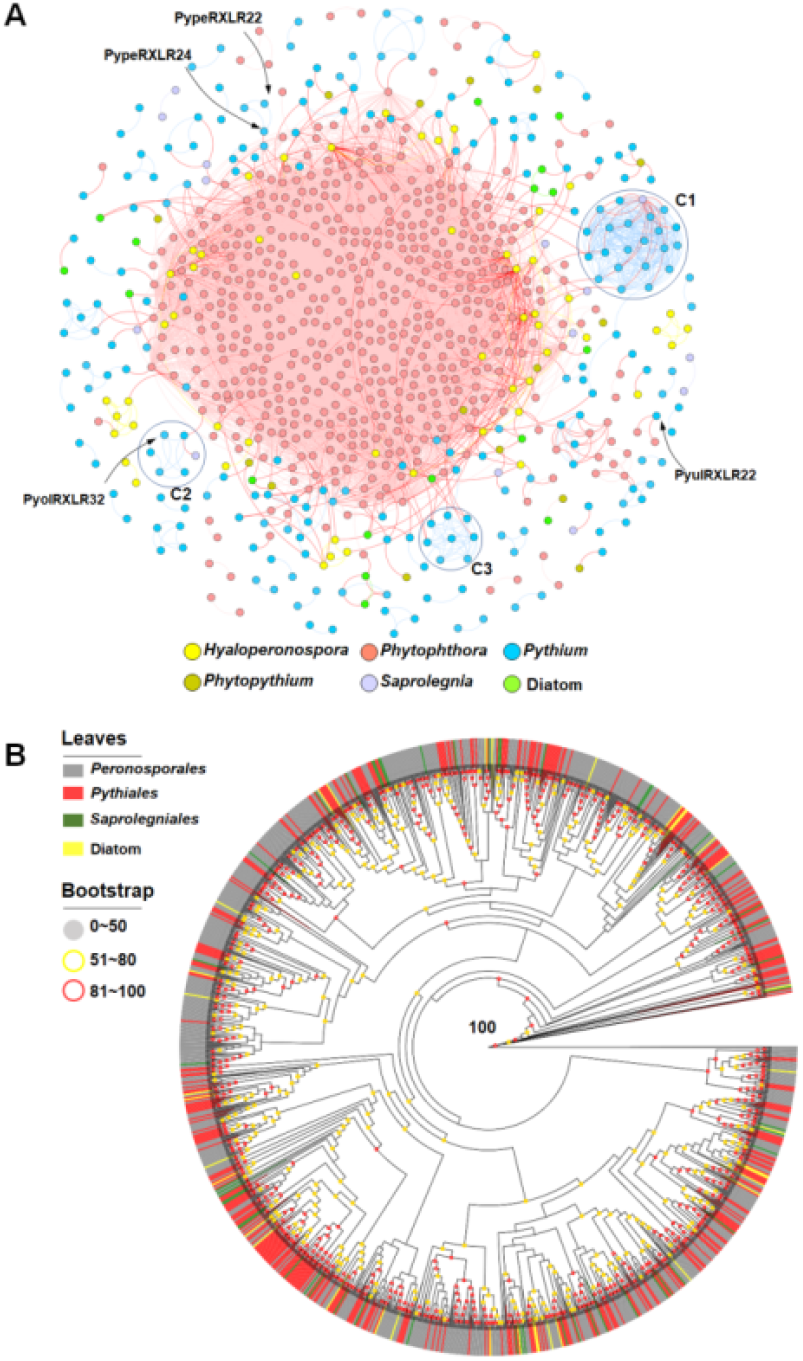
The evolutionary patterns of oomycetes RXLRs. (A). The relatedness network of putative RXLR effectors. Each spot represents an RXLR. Similar RXLRs (*E*-value<1) are joined by edges. Spot colors indicate their genera. Three relatively independent clusters were marked as C1, C2 and C3, respectively. Almost all effectors in these clusters are from *Pythium*. Four RXLRs with effector activities verified in this study are labeled using arrows. (B) The maximum-likelihood phylogenetic tree of the RXLR-dEER motifs in RXLR effectors. Grey, yellow and red circles in the nodes indicate bootstrap values of 0–50, 51–80 and 80–100, respectively. Leaves of the tree are colored base on the families of their indicated species.

Conserved RXLRs may play critical effector roles in pathogen virulence [10]. Analysis of shared sequence identity (peptides with BLASTP hit *E*-value<1e-5 and identity>30%) revealed 67, 93 and 28 *Pythium* RXLRs being intraspecies, intragenus and intergenus conserved, respectively (S5 Table), suggesting that *Pythium* RXLRs share a common ancestor with those from other genera and were only slightly expanded after species divergence. These conserved RXLRs may be essential effectors for pathogens and deserve more attention in future study.

Phylogenetic analysis was performed to further elucidate the evolutionary relationships among oomycete RXLRs as well as RXLR-like proteins from diatoms. Due to the host-pathogen arms race, RXLR C-terminals (after the dEER motif) are much more divergent [12] than their relatively conserved N-terminals [13,29]. Therefore, when building the phylogenetic tree, C-terminal and SP regions were discarded to reflect RXLR evolution relatedness with minimal interferences from host coevolution and secretory signal sequences. Except for PyolRXLR33 and HyarRXLR1, all other 1,111 RXLRs formed a huge clade (bootstrap value=100), indicating that they belong to a single superfamily. Using bootstrap value>80 as the criteria, 972 RXLRs can be divided into 105 subfamilies with member sizes ranging from 2 to 112 (S6 Table). No significant divergence was detected among *Peronosparales*, *Pythiales*, *Saprolegniales* or diatom RXLRs (Fig 2B). 39 and 6 subfamilies contain 119 and 28 RXLRs exclusively from *Peronosparales* and *Pythiales* species, respectively. All other subfamilies harbor RXLRs from at least two different genera. Taken together, the phylogeny results suggest a common ancestor shared by almost all RXLR and RXLR-like proteins with species-specific expansion much less detected in *Pythium* than in *Phytophthora*.

### Rapid evolution of *Pythium* RXLRs

Highly divergent *Phytophthora* and *Hyaloperonospor*a RXLRs are one of the most rapidly evolving portions in their proteomes [12,13]. Similarly, *Pythium* RXLRs also share low identities in our study. On average, only 37% identity was detected for all closest RXLR ortholog pairs within the *Pythium* genus. In contrast, we randomly selected 300 *Pythium* proteins and found a much higher average sequence identity (67%) associated with their closest ortholog pairs (Fig 3A). In our study, 233 (65%) *Pythium* RXLRs failed to form any ortholog pairs whereas only 10 out of 300 randomly selected *Pythium* proteins lack orthologs (Fig 3A). Furthermore, 236 (66%) *Pythium* RXLRs showed higher than 50% of sequence divergence (Fig 3A).

**Fig 3.**
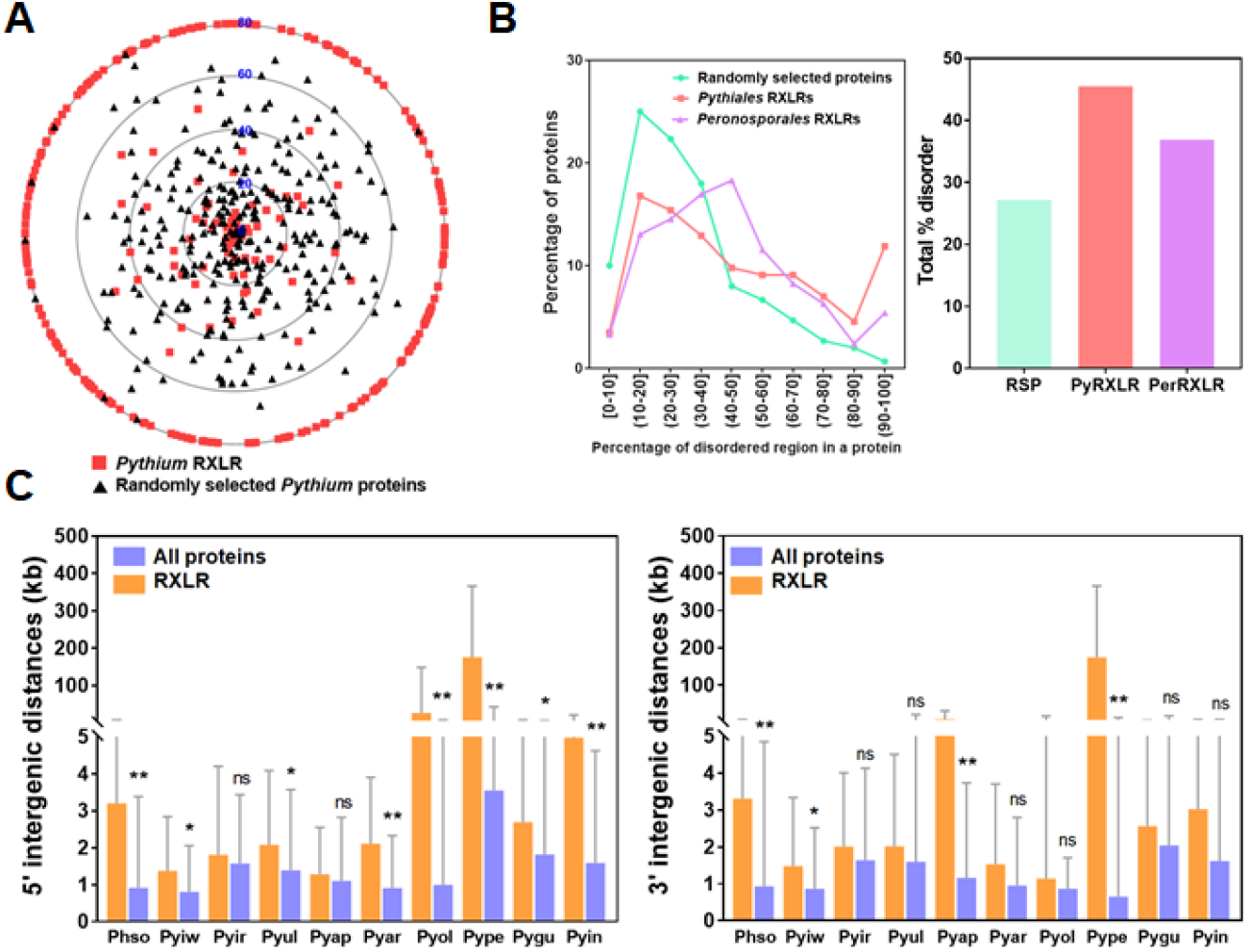
Evolutionary features of *Pythium* RXLRs. **(A) Sequence divergence of *Pythium* RXLRs**. *Pythium* RXLRs were compared against 300 randomly selected *Pythium* proteins regarding the sequence identities of their closest ortholog pairs within the *Pythium* genus. Radiuses range from 0 (center) to 80 (outer circle) represent 100% to 20% (or less) sequence identities sequentially. Proteins at the same identity level are randomly distributed along their corresponding circle. **(B) Disorder content in***Peronosporales* **and***Pythiales* **RXLRs.** The line plot shows the percentages of proteins containing different percentages of disordered regions. The bar plot shows total disorder percentages in *Peronosporales*, *Pythiales* and randomly selected proteins. RSP, Randomly selected proteins; PyRXLR, *Pythiales* RXLRs; PerRXLR, *Peronosporales* RXLRs. **(C) 5’ and 3’ end intergenic distances of** *P. sojae* **and***Pythium RXLRs*. The two bar plots show 5’ (left) and 3’ (right) end intergenic distances of indicated *RXLR* genes, respectively. (*, *P*<0.05; **, *P*<0.001, Student’s *t* test). Phso*, P. sojae*; Pyiw, *P. iwayamai*; Pyir, *P. irregulare*; Pyul, *P. ultimum*; Pyap, *P. aphanidermatum*; Pyar, *P. arrhenomanes*; Pyol, *P. oligandrum*; Pype, *P. periplocum*; Pygu*, P. guiyangense*; Pyin, *P. insidiosum*.

### Disorder content is abundant in *Pythium* RXLRs

Disorder content is abundant in *Phytophthora* RXLRs which contributes to their effector activities [16]. Our analysis demonstrated the abundance of disorder content in *Pythium* RXLRs as well. The mean percentages of disorder region in randomly selected *Pythium* proteins, *Pythium* RXLRs and *P. sojae* RXLRs are 27.17%, 45.48% and 36.89%, respectively (Fig 3B). *Pythium* RXLRs shows obviously abundant disorder content which is even higher than that of *P. sojae*. Over 64% *Pythium* RXLRs have 30% or more disordered residues, which can be found in only 43.67% randomly selected *Pythium* proteins (Fig 3B).

### *Pythium* RXLRs are predominantly located in gene-sparse regions

*Phytophthora RXLR* genes are preferably located in gene-sparse regions [15]. In *P. sojae*, both the 5’ and the 3’ end average intergenic distances of *RXLRs* are significantly longer than those of the complete *P. sojae* genes (*P*<0.05, Student’s *t* test). The mean 5’ end intergenic distance of *RXLRs* is 3.20 kb with the whole-genome mean being 0.91 kb. *RXLRs*’ average 3’ end intergenic distance of 3.31 kb is also much longer than that of the complete *P. sojae* genes (0.92 kb) (Fig 3C). Both observations are similar to the previous report [15].

Similar gene location preference was detected for *Pythium RXLRs* in our study. Compared with the complete gene sets of their corresponding species, *RXLRs* of all 9 examined *Pythium* species have longer average intergenic distances at both 5’ and 3’ ends (Fig 3C). For the 5’ and the 3’ end average intergenic distances, the differences are statistically significant (*P*<0.05, Student’s *t* test) in 7 and 3 *Pythium* species, respectively (Fig 3C). These results indicate that *Pythium RXLRs* are predominantly located in gene-sparse regions, which is similar to that of the *Phytophthora RXLRs*.

### Genome rearrangement contribute to the emergence of novel *RXLR* genes in *P. guiyangense*

*P. guiyangense* has a larger RXLR repertoire than all other *Pythium* species examined (Fig 1B). Our previous synteny analysis showed that *P. guiyangense* has a hybrid genome derived from two distinct parental species, which leads to a nearly two-fold expansion of its predicted genes when compared with other *Pythium* species. Consequently, most homology gene pairs in *P. guiyangense* show high degrees of collinearity [24]. However, we only found 4 *RXLRs* from different scaffolds showing collinearity, indicating that additional mechanisms may also contribute to the expansion of *RXLR* genes in *P. guiyangense*. We noticed that 29 *PyguRXLRs* form several clusters (<40 kb in size) with each cluster located in the rearrangement region of a same contig. Similar phenomenon was not observed in other *Pythium* species (S2 Fig). For example, *PyguRXLR6/7/8* (Fig 4A) and *PyguRXLR32/33/34* (Fig 4B) are clustered in the genome rearrangement regions in contigs 2 and 9, respectively. Furthermore, 22 clustered RXLRs show high similarities in N-terminals but have less conserved or even highly divergent C-terminals within each cluster. For example, PyguRXLR6/7/8 share a conserved N-terminal but their C-terminals are highly divergent (Fig 4C). Similar pattern was observed in PyguRXLR32/33 versus PyguRXLR34 (Fig 4D). Taken together, these results suggest that both gene duplications and genome rearrangements contribute to the expansion and the emergence of novel *RXLRs* in *P. guiyangense*.

**Fig 4.**
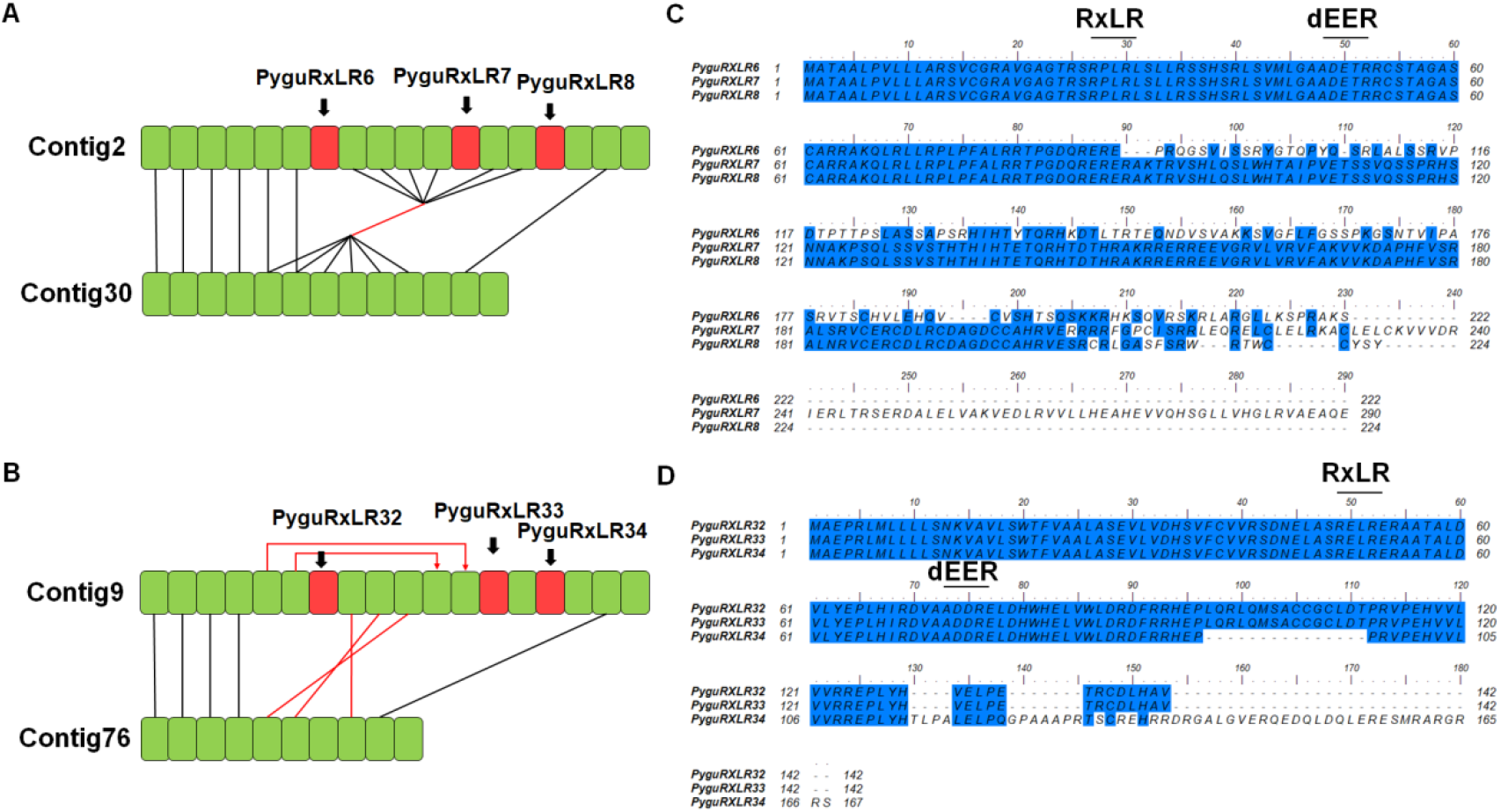
Synteny analysis and protein sequence alignments of clustered RXLRs in *P. guiyangense*. (A and B) Synteny analysis of clustered *P. guiyangense RXLRs*. Red blocks indicate clustered *PyguRXLRs* described in the text. Green blocks indicate genes flanking to these *PyguRXLRs*. Homologous genes with similar and opposite genome orientations are joined with black and edges, respectively. (C and D) Protein sequence alignments of two RXLR clusters. Identical or similar residues are colored in blue. The RXLR-dEER motifs are labeled in the alignments.

### Most predicted *P. ultimum RXLRs* are truly transcribed genes

To further elucidate whether these predicted *Pythium RXLRs* are truly transcribed genes, an RNA-seq analysis was carried out to examine the transcript accumulations of 40 *RXLR* candidates in *P. ultimum* during mycelium and 3, 6, 12, 24, 36 h post host-infection stages. As one of the most important oomycetous plant pathogens with a high-quality assembled genome sequence available [21,30], *P. ultimum* was selected as the investigation target. Except for 2 candidates (PyulRXLR25/26) whose RPKM (reads per kilobase transcript length per million reads mapped) values were consistently below 1 at all stages, transcripts of the remaining 38 predicted *RXLRs* can be detected at one or more stages (Fig 5A), suggesting that most *P. ultimum RXLRs* predicted in our study are truly transcribed genes.

**Fig 5.**
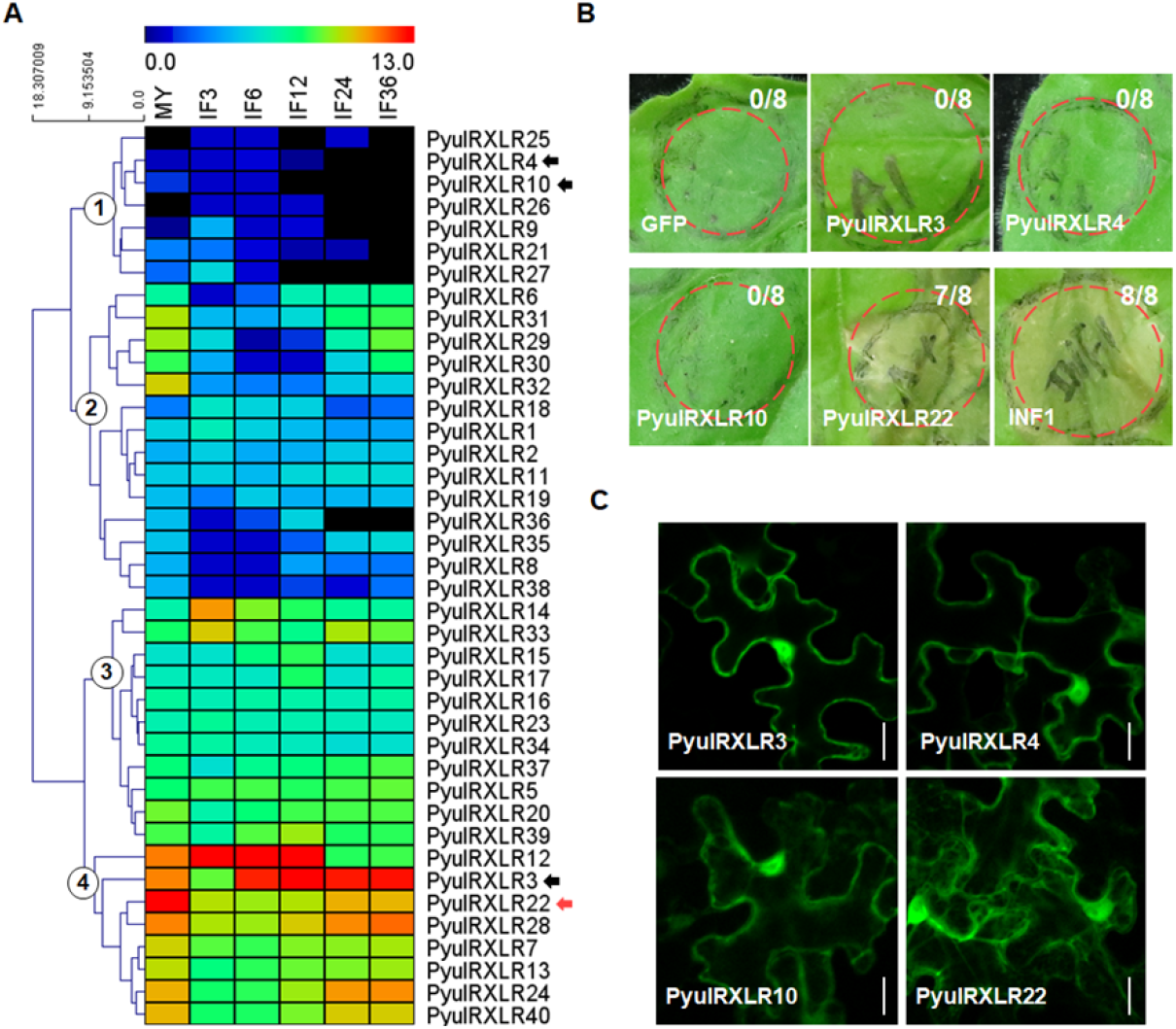
Transcription patterns of *P. ultimum RXLRs* and the effector activity of PyulRXLR22. (A) Transcription patterns of *P. ultimum RXLR* candidates. The heat map shows gene transcription patterns at the mycelia (MY) stage and stages of 3, 6, 12, 24 and 36 h post-inoculation of soybean leaves. Color bars represent log2 of gene RPKM values, ranging from dark blue (0) to red (13). Black bars indicate no expression detected. RXLRs selected for the effector activity assay are marked with arrows with red arrow indicating ability of inducing cell death in *N. benthamiana*. Based on transcript accumulation levels and patterns, *P. ultimum RXLRs* are divided into 4 clusters using HCL methods and the MeV software. Clusters are marked with circled numbers (1 to 4). (B) PyulRXLR22 induced cell death in *N. benthamiana*. *N. benthamiana* leaves were transformed with the indicated constructs by agro-infiltration. Dashed circles indicate infiltration sites and the ratios are sites with necrotic lesion versus total infiltration sites. PyulRXLR3/4/10 did not induce cell death. GFP and INF1 were used as negative and positive controls, respectively. (C) Subcellular localization of PyulRXLRs. Green fluorescence indicates the localizations of PyulRXLRs in *N. benthamiana* epidermal cells. Photographs were taken at 48 hpi. Bar =20 μm.

*P. ultimum RXLR* genes showed stage-specific expression patterns. They were assigned to four gene expression clusters (Fig 5A) via hierarchical clustering (HCL). Cluster 1 contains 2 putative pseudogenes and 5 *RXLRs* exhibiting zero to low expression. Despite their low transcript accumulations, all *RXLRs* in this cluster showed elevated expression during early infection stages. The 14 *RXLRs* in Cluster 2 exhibited elevated transcript accumulations in both mycelium and late infection stages. The expression of 11 Cluster 3 *RXLRs* were largely constant across all the stages with slightly more transcripts detected at 3/6 hours post infection. The 8 Cluster 4 *RXLRs* showed a similar expression pattern as that of the *RXLRs* in Cluster 2 but with much higher RPKM values. These results revealed major transcription shifts of *PyulRXLRs* at mycelium and different host infection stages.

### PyulRXLR22 induces cell death in *Nicotiana benthamiana*

*Phytophthora* RXLRs suppress PAMP-induced plant cell death. Their ectopic expression can trigger cell death in plants [31]. To evaluate the effector activity of predicted PyulRXLRs, we selected 4 candidates (PyulRXLR3/4/10/22, 2 from Cluster 1 and 2 from Cluster 4) for functional characterizations (Fig 5A). When co-expressed with *INF1* [32] in *N. benthamiana* leaves, none of these four RXLRs can suppress cell death triggered by the *P. infestans* PAMP INF1 (S3 Fig). When evaluating their ability to trigger host cell death by ectopic expression, only PyulRXLR22 induced dramatic cell death in *N. benthamiana* leaves (Fig 5B). *PyulRXLR22* expression is highly upregulated at late infection stages (Fig 5A). Fused with a GFP tag at N-terminal, the four PyulRXLRs were transiently expressed in *N. benthamiana* epidermal cells to reveal their subcellular localizations. PyulRXLR22 is located in endoplasmic reticulum (ER), plasma membrane and the nucleus (Fig 5C). In contrast, PyulRXLR3/4/10 were found in plasma membrane and the nucleus but not in ER (Fig 5C). Bioinformatic analysis showed that PyulRXLR22 has two homologs in *P. ramorum* (BLASTP *E*-value<1) (Fig 2A). The demonstrated effector activity makes PyulRXLR22 and its homologs promising targets for further investigation.

### RXLRs from biocontrol *Pythium* species induce plant defense in *N. benthamiana*

*P. oligandrum* and *P. periplocum* are two unique *Pythium* species that are widely used for controlling soilborne plant diseases. They can infect pathogens and induce plant defense reactions as well [25,26,27]. To elucidate whether their RXLRs participate in this biocontrol process, all 73 RXLRs of these two species were cloned and expressed in *N. benthamiana* leaves individually to test their ability of triggering cell death. PyolRXLR32, PypeRXLR22 and PypeRXLR24 triggered cell death at 7 days post infiltration (dpi) (Fig 6A). All other 70 RXLRs lack this ability. PyolRXLR32 is located in plasma membrane and the nucleus. Both PypeRXLR22 and PypeRXLR24 are located in plasma and nuclear membranes (S4 Fig).

**Fig 6.**
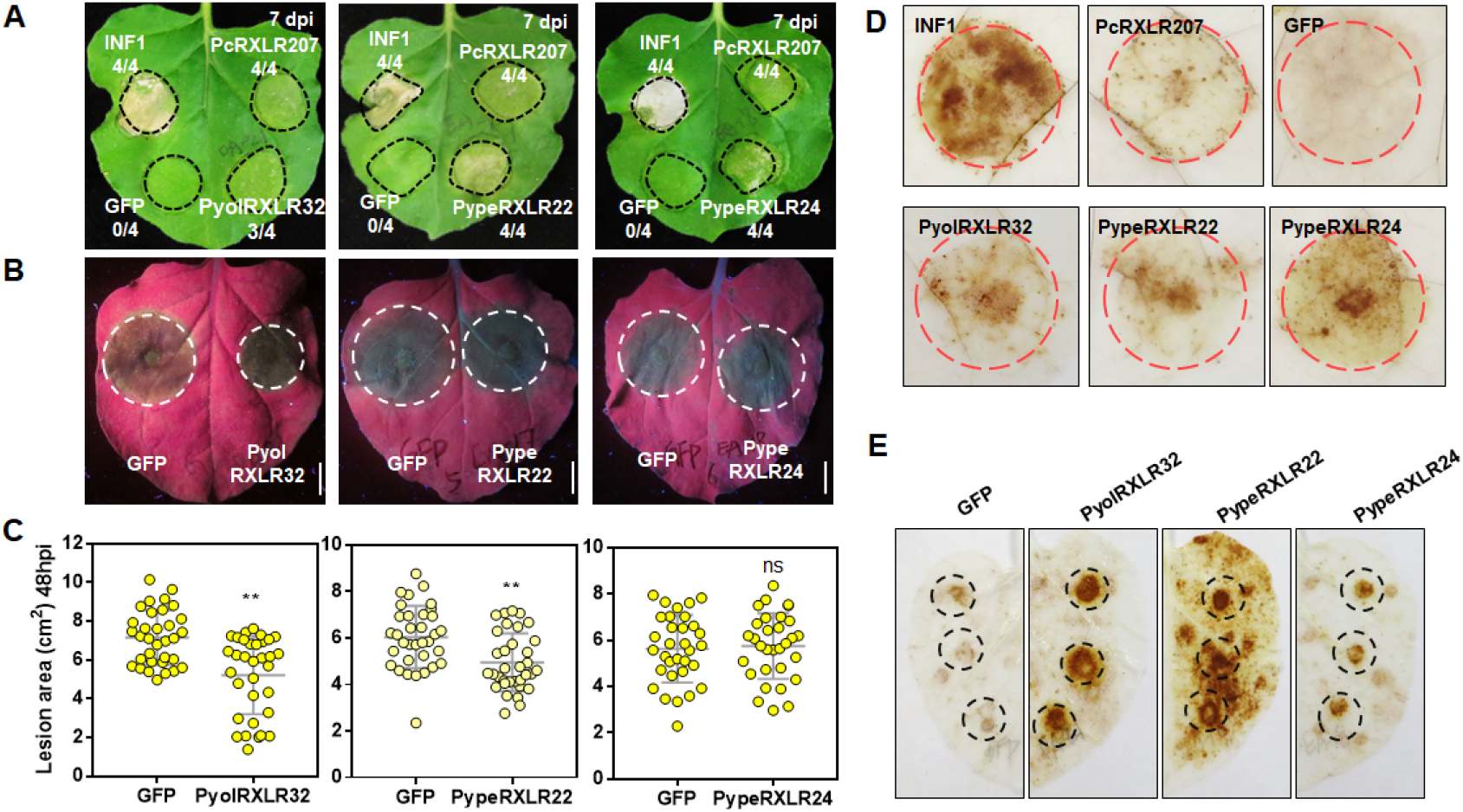
RXLRs from *P. oligandrum* and *P. periplocum* induce plant defense response. (A) Expression of PyolRXLR32, PypeRXLR22 or PypeRXLR24 induce cell death in *N. benthamiana* leaves. *N. benthamiana* leaves were transformed with the indicated constructs by agro-infiltration. Dashed circles indicate infiltration sites and the ratios are sites with necrotic lesion versus total infiltration sites. INF1 and the RXLR effector RXLR207 from *P. capsici* were used as positive controls. Empty vector expressing GFP was used as a negative control. Same controls were also used in the H_2_O_2_ accumulation assay. (B and C) *P. capsici* infection can be reduced by PyolRXLR32 and PypeRXLR22, but not PypeRXLR24. 24 h after infiltration, equal amount of mycelium was inoculated onto each infiltrated area expressing *PyolRXLR32*, PypeRXLR22, or PypeRXLR24. Photographs were taken under UV light at 48 hpi and lesion areas were measured at the same time. Lesion area data shown in the dot plots were collected from three biological replicates. At least five leaves were inoculated for each replicate (**, *P*<0.001, Student’s *t* test). Bar=1cm. (D) H_2_O_2_ accumulation in *N. benthamiana* leaves expressing PyolRXLR32, PypeRXLR22 or PypeRXLR24. DAB staining was performed 48 hpi. Red dashed circles indicate infiltration sites. (E) H_2_O_2_ accumulation in *P. capsica*-inoculated *N. benthamiana* leaves expressing PyolRXLR32, PypeRXLR22, or PypeRXLR24. *P. capsica* was inoculated 24 h after infiltration. DAB staining was performed at 12 hpi. Black dashed circles indicate inoculation sites.

To investigate whether the three RXLR effectors can induce plant defense, we infiltrated *N. benthamiana* leaves with *Agrobacterium tumefaciens* harboring a *GFP* or indicated *RXLR* fusion construct and then inoculated the leaves with *P. capsici*. Compared with the GFP control, both PyolRXLR32 and PypeRXLR22 significantly reduced the lesion area caused by *P. capsici* whereas PypeRXLR24 did not have noticeable effect on lesion control (Fig 6B and 6C).

Reactive oxygen species (ROS) play a central role in plant immunity with ROS burst being a defense response indicator [33]. Our DAB staining assay showed that, compared with the GFP control, ectopic expression of any of the three RXLRs induced ROS burst in *N. benthamiana* leaves (Fig 6D). Furthermore, all three RXLRs can enhance ROS burst triggered by *P. capsici* inoculation (Fig 6E).

Interestingly, PyolRXLR32 is both intragenus and intergenus conserved (S5 Fig). Its *P. infestans* homolog (BLASTP *E*-value=2e-115), previously reported as SFI4 (Suppressor of early Flg22-induced Immune response 4), can inhibit host immunity and promote infection [34], which is opposite to PyolRXLR32’s positive role in triggering cell death and defense response. Likewise, although PypeRXLR21/22/24 share high sequence similarity (S6 Fig), PyolRXLR21 cannot trigger host cell death as PyolRXLR22/24 did. These results indicate that slight sequence difference may lead to diverse or even contrary functions of homologous RXLR effectors.

## Discussion

In this study, we performed a *de novo* identification of RXLR effector genes from the genomes of nine *Pythium* species and analyzed their evolutionary features. Selected *Pythium* RXLR effectors were functionally verified. Despite the stringent prediction method we adopted, RXLRs can be found in a wide range of oomycetes including the *Pythium* species, and are almost absent outside *Stramenopiles*. Our phylogenetic analysis indicates the existence of a common ancestor for RXLRs with all oomycetous RXLRs clustered into a single superfamily. *Pythium* and *Phytophthora RXLR* genes share several similar features including rapid evolution, abundant disorder content and the preference of locating in gene-sparse regions. In particular, *P. guiyangense* harbors a much larger RXLR reservoir than other *Pythium* species due to frequent gene duplication and genome rearrangement events. Using *P. ultimum* as a model, most of its predicted *RXLRs* were demonstrated to be truly transcribed genes that cluster into four distinct expression patterns. The effector activity of PyulRXLRs22 was functionally verified in this study. Finally, we identified three functional RXLR effectors from two biocontrol *Pythium* species.

Although no *Pythium* RXLRs has been identified before [21,22], there are clues indicating their existence. Several RXLR effectors have been experimentally validated in species phylogenetically close to *Pythium*, including *A. candida*, *A. laibachii* and *S. parasitica* [17][18] [19]. The necrotrophic infection type of *Pythium* pathogens also match the observations that many RXLR effectors can induce necrosis. Largely due to the stringent definition of the dEER motif, existing genome annotation models may be inadequate for *RXLR* gene prediction in *Pythium*. Hence, we developed a modified regex model to allow the search of degenerate dEER motifs. Previously reported effectors were adopted as references and all 6 possible ORFs were retrieved from the whole genomes. To reduce false-positive rates, we omitted the BLASTP step following RXLR searching. By adopting our modified regex model, we predicted 359 new RXLR proteins in nine *Pythium* species as well as re-identification of RXLRs from *H. arabidopsidis*, *P. ramorum* and *P. sojae*. The high accuracy of our new model is demonstrated by the observations that very few RXLRs was predicted outside *Stramenopiles* and all re-identified RXLRs are subsets of previously known RXLRs. Notably, our detection of RXLR-like proteins in diatoms suggests the origin of oomycetous RXLRs from an early ancestor of the *Stramenopiles* lineage. *Pythium* and other oomycetous RXLRs form a superfamily in our phylogenetic analysis, which also support the common-ancestor hypothesis of RXLR evolution.

Despite the high divergence of RXLR sequences, oomycetous RXLRs identified in our study are close enough to form a huge relatedness network with *Phytophthora* RXLRs exhibiting higher overall sequence similarities. There are also RXLR clusters predominantly associated with *Pythium* or *Hyaloperonospora* species, suggesting the occurrence of independent evolution in these effectors. *Phytophthora* and *Pythium* RXLRs share several common features.

*Phytophthora* RXLR effectors promote virulence by manipulating host defense processes. Thus, they undergo rapid sequence evolution to escape the surveillance of plant immune system. In our study, this fast evolution pattern was also observed in *Pythium* RXLRs, which indicate their participation in host pathogenesis. As a consequence of effector evolution, abundant disorder regions exist in *Phytophthora* RXLRs [16]. These disorder regions are even more prevalent in *Pythium* RXLRs. *Phytophthora* genomes feature a so-called ‘two-speed’ structure of gene-dense and gene-sparse regions [35], with *RXLRs* enriched in the more dynamic gene-sparse regions. Gene-sparse regions are the heat sites of genome rearrangement and tend to accelerate pathogen evolution [36]. In *Pythium* species, we also found that both 5’ and 3’ end average intergenic distances of *RXLRs* are significantly longer than the whole genome averages, which demonstrates the similar location preference of *Pythium RXLRs* in the gene-sparse regions.

Due to their highly divergent C-terminals, few RXLR paralogs can be found in *Pythium* species except for *P. guiyangense*. We characterized the *P. guiyangense* genome as hybrid and identified a large set of *RXLR* genes. *P. guiyangense RXLRs* share low degrees of synteny with *PyguRXLR* paralogs often clustered in the same contig and located in genome rearrangement regions. Furthermore, physically close paralog pairs generally share a conserved N-terminal but have less conserved or even highly divergent C-terminals. Our results indicate that recently happend gene duplication and genome rearrangement events shape the evolution theme of *P. guiyangense RXLR*s. Previous research suggests that the duplications of genes encoding virulence-associated effectors can facilitate pathogen adjustment towards a host jump [37]. In the case of *P. guiyangense*, the duplication and rearrangement events detected may generate novel *RXLR* genes which can help its adaption against insects. *RXLR* genes also clustered closely (<100kb) in *P. ramorum* [12], suggesting that *RXLR* gene cluster may be a common host adaptation strategy shared by oomycete species.

Our RNA-seq analysis on *P. ultimum* demonstrated that most predicted *RXLRs* are transcribed genes, which further verified the reliability of our redesigned RXLR annotation method. Meanwhile, these *RXLRs* exhibit distinct expression patterns during infection. RXLR expression stages are associated with their specific functions. In *P. sojae*, early-induced RXLRs inhibit host cell death triggered by PAMPs, while necrosis is caused by late-induced RXLR effectors [31]. Another *Phytophthora* pathogen, *P. capsici*, secretes a RXLR effector to induce host defense and cell death, and trigger the biotroph-to-necrotroph transition [9]. Since most *Pythium* pathogens are necrotrophic, their RXLRs may trigger host cell death instead of acting as cell death repressors. Consistent with this expectation, none of our tested PyulRXLRs can suppress host cell death induced by INF1. Instead, ectopic expression of PyulRXLR22 triggers necrosis. The function diversity of RXLRs in *Phytophthora* and *Pythium* species may result from their different nutrient acquisition strategies during infection.

*P. oligandrum* and *P. periplocum* are two biocontrol agents with their defense-inducing mechanisms largely unknown. Previous reports suggest that plant defense may be induced by Microbe-Associated Molecular Patterns (MAMPs) from these two *Pythium* species [38]. In this study, we identified three RXLR effectors from them and demonstrated the plant defense induction ability of these RXLRs. This finding unveils the participation of cytoplasmic effectors in triggering host defense reactions. Since the *P. oligandrum* hyphae can penetrate into the root tissues, its RXLR effectors may be secreted to host cells via a similar mechanism detected in *Phytophthora* pathogens.

In conclusion, we for the first time identified RXLR effectors in *Pythium* species at the genome-wide scale. Their evolutionary features, sequence characteristics and expression patterns were analyzed in detail. The effector activities of selected *Pythium* RXLRs were experimentally verified, including their diversified functions in inducing cell death and host defense. This study expands our current understanding of oomycetous RXLR effectors to the *Pythium* genus. Resources generated in this work facilitate future in-depth investigations on the interactions between these novel *Pythium* RXLRs and their diversified hosts.

## Materials and methods

### Data sets

The genome sequences of *P. ramorum* and *P. sojae* [4] were obtained from the Joint Genome Institute website (http://genome.jgi-psf.org). The genome sequences of *H. arabidopsidis* [28], *Ph. vexans* [22], *P. aphanidermatum* [22], *P. arrhenomanes* [22], *P. irregulare* [22], *P. iwayamai* [22], *P. oligandrum* [39], *P. periplocum* [39], *P. ultimum* [21], *P insidiosum* [23], *P. guiyangense* [24], *S. parasitica* [40], *P. tricornutum* [41], *T. pseudonana* [42], *S. cerevisiae* [43], *S. sclerotiorum* [44], *U. maydis* [45], *P. graminis* [46], *B. cinerea* [47], *C. elegans* [48], *M. hapla* [49], *E. coli* [50], *E. amylovora* [51] and *Ca* L. asiaticus [52] were obtained from the National Center for Biotechnology Information (NCBI, https://www.ncbi.nlm.nih.gov/). Detailed genomic information of all species is listed in S7 Table.

### Identification of RXLR effectors

For RXLR effector identification, all open reading frames (ORFs) encoding proteins longer than 50 amino acids were retrieved. Signal peptides (SP) were predicted using SignalP v3 with the criteria that cleavage sites are located between residues 10 and 40 and HMM prob>=0.9 [53]. Transmembrane regions (TMs) were predicted using TMHMM v2.0 [54]. Sequences containing SP but lacking TM were retained and screened for RXLRs with a modified regex model described as SP.{1,40}R.LR.{1,40}([ED].[ED][KR]|[ED][ED].{0,3}KR]). All matched protein sequences were predicted as RXLR effectors. For redundant RXLRs caused by a set of alternative overlapping ORFs within the same reading frame, only the single ORF encoding the longest RXLR was selected.

### Phylogenetic analysis

For phylogenetic analysis, SPs and C-terminals after the dEER motif were removed from RXLR sequences. The truncated sequences were aligned using MUSCLE [55] and then manually edited. Maximum-likelihood (ML) trees were constructed using the IQ-TREE software with the parameter of -bb 1000 [56]. VT+F+R6 was selected as the best-fit model for this assay by using the ModelFinder tool in IQ-TREE. The phylogenetic tree was showed using EvolView v3 [57].

### Disorder region prediction and synteny analysis

Disorder region prediction was performed by using ESpritz (http://protein.bio.unipd.it/espritz/) with the X-ray prediction type and the best-sw threshold [58]. Synteny analysis was conducted using MCScanX [59].

### Vector construction

For transient expression in *N. benthamiana*, *PyulRXLRs, PyolRXLRs* and *PypeRXLRs* (lacking the SP-encoding region) were amplified from their corresponding genomic DNA and inserted into the pBinGFP2 vector [60]. Primers used for PCR amplification are listed in S8 Table.

### Plant materials and agroinfiltration

*N. benthamiana* plants were maintained in a greenhouse with an environmental temperature of 25°C under a 16-h/8-h light/dark photoperiod. The *A. tumefaciens* strain GV3101 was previously stored in our lab. Before infiltration, *A. tumefaciens* clones harboring indicated constructs were cultured at 28°C, 200 rpm for 48 h to reach the appropriate concentration. The *Agrobacterium* cultures were then resuspended in 10 mM MgCl_2_ to an appropriate optical density (OD). For the necrosis induction assay, *A. tumefaciens* cells were infiltrated into the 6-week-old leaves of *N. benthamiana* at a final OD of 0.2 at 600 nm for each construct. For the cell death suppression assay, *A. tumefaciens* cells expressing the indicated RXLR effectors (OD_600_=0.3) were infiltrated into *N. benthamiana* leaves one day before the infiltration of *A. tumefaciens* expressing INF1 (OD_600_=0.2). Symptom development was examined 7 days after infiltration.

### Oomycete culture conditions and the inoculation assay

The *P. oligandrum* strain CBS 530.74 and the *P. periplocum* strain CBS 532.74 were kindly provided by Dr. Laura J. Grenville-Briggs at the Swedish University of Agricultural Sciences and were stored at 25 °C on 10% (v/v) V8 juice medium in the dark. *P. ultimum* and the *P. capsici* strain LT263 were previously stored in our lab and grown at 25 °C on the V8 medium in the dark. For *P. capsici* mycelium inoculation, 7-mm disks of 4-day growth medium were inoculated on *N. benthamiana* leaves 24 h after infiltration. Inoculated leaves were photographed at 48 hpi under UV light. Lesion areas were measured at the indicated time points. This assay was independently repeated for at least three times. For the inoculation of *P. ultimum*, 4-mm plugs from the growing edge of a 2-day V8 agar culture were placed in the *N. benthamiana* leaves, and the samples were harvested at the indicated time points.

### Transcriptome sequencing and bioinformatic analysis

For mycelium samples, *P. ultimum* mycelium growing in V8 agar culture for 2 days were harvested. *P. ultimum* samples at different infection stages (3, 6, 12, 24, 36 hpi) were harvested using a 5 cm punch. Three biological replicates were performed and used to produce an independent pool. The quality of total RNA was evaluated by Agilent 2100 Bioanalyzer (Agilent Technologies, Santa Clara, CA, USA). Samples with OD_260/230_≥1.8 and OD_260/280_≥1.8 were used for RNA-seq. Raw reads were filtered using the FastQC software (https://www.bioinformatics.babraham.ac.uk/index.html) to remove poor-quality reads and adapters. Sequences with a Phred score above 20 were considered as clean reads. All clean reads were mapped to the *P. ultimum* reference genome using Hisat2 with the parameters of --min-intronlen 30 --max-intronlen 5000 –dta [61]. SAMtools was used to convert and sort SAM files [62]. Based on the lengths of each gene and the counts of uniquely mapped reads, gene expression levels were normalized as RPKM (reads per kilobase transcript length per million reads mapped) using the Stringtie tool [61]. Heat maps and clustering (using hierarchical clustering methods) of gene expression patterns were performed using the MeV software [63].

### DAB staining

*N. benthamiana* leaves with ectopic expression of indicated RXLR effectors were detached at 48 hpi and then stained with 1 mg/ml DAB solution for 8 h in the dark. After destaining with ethanol, leaves were examined under light microscopy. *P. capsica*-inoculated *N. benthamiana* leaves were stained with 1 mg/ml DAB solution for 8 h in the dark at 12 hpi, and then destained with ethanol before photographing.

## Supporting information

**S1 Fig. Sequence alignment of** *Pythium* **RXLRs.** Signal peptide and the RXLR-dEER motifs are labeled above their corresponding regions. Arg and Lys residues in the RXLR-dEER motifs are colored in blue. Glu and Asp residues are colored in red.

**S2 Fig. Numbers of clustered** *RXLR* **effectors in each species.** *RXLR* pairs with distances less than 40 kb in the same scaffold are showed. Hyar, *H. arabidopsidis*; Phra, *P. ramorum*; Phso*, P. sojae*; Phve, *P. vexans*; Pyiw, *P. iwayamai*; Pyir, *P. irregulare*; Pyul, *P. ultimum*; Pyap, *P. aphanidermatum*; Pyar, *P. arrhenomanes*; Pyol, *P. oligandrum*; Pype, *P. periplocum*; Pygu*, P. guiyangense*; Pyin, *P. insidiosum*; Sapa, *S parasitica*; Thps, *T. pseudonana*; Phtr, *P. tricornutum*.

**S3 Fig. PyulRXLRs do not inhibit cell death triggered by INF1**. *A. tumefaciens* harboring indicated constructs were infiltrated into *N. benthamiana* leaves 24 h before the infiltrations of INF1. Photos were taken 4 days after infiltration.

**S4 Fig. Subcellular localization of PyolRXLR32, PypeRXLR22 and PyolRXLR24.** Indicated constructs were expressed in *N. benthamiana* epidermal cells to show protein localizations. Images were taken at 48 hpi. Bar=20μm.

**S5 Fig. Sequence alignment of PyolRXLR32 and its homologs.** The RXLR-dEER motifs were labeled above their corresponding regions.

**S6 Fig. Sequence alignment of PyolRXLR21, 22 and 24**. The RXLR-dEER motifs were labeled above their corresponding sites.

**S1 Table.** Overall counts of predicted RXLR effectors per species.

**S2 Table.** Known functional RXLR with degernerate dEER motif

**S3 Table.** Enrichment assay result comparing *B. cinerea* to the other species

**S4 Table.** Permutation test of RXLR motif in each species

**S5 Table.** Conserved RXLR effectors

**S6 Table.** RXLRs summary

**S7 Table.** Genomic information used in this study

**S8 Table.** Primers used in this study

**S9 Table.** RPKM value of *PyRXLRs* during different stages

